# A minimal thermodynamic theory for re-entrant liquid-liquid phase separation regulated by small molecules

**DOI:** 10.64898/2026.06.12.731829

**Authors:** Achal Jadhav, Pushpita Ghosh

**Affiliations:** School of Chemistry, Indian Institute of Science Education and Research, Thiruvananthapuram, Kerala 695551, India; Center for High-Performance Computing, Indian Institute of Science Education and Research Thiruvananthapuram, Kerala 695551, India

## Abstract

Small molecules regulate biomolecular condensates in a biphasic manner, promoting liquid–liquid phase separation (LLPS) at low concentrations while suppressing it at higher concentrations. Despite increasing experimental evidence for such re-entrant behavior, a unified physical description remains lacking. Here, we identify a minimal thermodynamic mechanism for re-entrant LLPS by coupling Cahn–Hilliard dynamics to a concentration-dependent Flory interaction parameter containing competing LLPS-promoting and inhibitory contributions. The resulting model reproduces experimentally observed nonmonotonic condensate formation in Tau–tannic acid and TDP-43–bis-ANS systems, including the concentration-dependent emergence and dissolution of protein-rich domains. Spinodal analysis reveals finite concentration windows for phase instability and demonstrates that re-entrant mixing is encoded directly in the free-energy landscape. The framework further captures morphology transitions and diffusive coarsening within the phase-separated regime. These results establish a general mesoscale description of chemically regulated condensates and provide design principles for controlling phase separation through small-molecule modulators.

## Introduction

Liquid-liquid phase separation (LLPS) is a key physical mechanism by which cells form biomolecular condensates that spatially organize biochemical components in the absence of membranes.^1–5^ Recent years have witnessed significant advances in understanding the physical principles governing condensate formation through both passive equilibrium processes and active, energy-driven mechanisms.^6,7^ The physics of droplet formation, regulation, and material properties in biological cells continues to be an active area of theoretical and experimental investigation. ^8–11^ These condensates include cytoplasmic structures such as P-bodies and stress granules,^12,13^ as well as nuclear bodies including nucleoli^14^ and Cajal bodies.^15^ Many intrinsically disordered proteins (IDPs), including TAR DNA-binding protein 43 (TDP-43)^16^ and Tau,^17^ undergo LLPS under physiological conditions to form dynamic condensates involved in cellular organization, signaling, and stress adaptation.

Recent studies have further shown that the stability and material properties of biomolecular condensates can be strongly modulated by metabolites and small molecules, which alter intermolecular interactions and condensate formation.^18–22^ Depending on concentration, such modulators may either promote LLPS through transient bridging interactions^23,24^ or suppress condensation through hydrotropic solvation and interaction screening. ^25–27^ Dysregulation of these processes is increasingly linked to pathological aggregation and neurodegenerative disease. For example, aberrant phase behavior of TDP-43 is associated with amyotrophic lateral sclerosis (ALS),^28^ while Tau aggregation is implicated in Alzheimer’s disease.^29,30^

Experiments have revealed that several small molecules regulate LLPS in a biphasic manner, promoting condensate formation at low concentrations while suppressing phase separation at higher concentrations.^31^ Such re-entrant behavior has been reported for TDP-43 in the presence of ATP or bis-ANS,^31,32^ Tau with tannic acid,^33^ FUS with ATP,^27^ and RNA-mediated polypeptide systems.^34,35^ These observations suggest that competing molecular interactions can alter condensate stability in qualitatively different ways across concentration regimes, giving rise to finite concentration windows of phase separation.

Despite these experimental advances, a unified physical description of biphasic condensate regulation remains lacking. Previous theoretical studies have primarily focused either on LLPS enhancement at low modulator concentrations using continuum reaction–diffusion descriptions,^36^ or on suppression of LLPS at high concentrations using molecular-dynamics simulations and mean-field approaches.^25,26,37^ Consequently, the minimal thermodynamic ingredients required to generate re-entrant phase behavior remain unclear. Beyond explaining existing experiments, identifying the minimal physical ingredients responsible for re-entrant LLPS is important for developing predictive theories of chemically regulated condensates. A minimal description can help distinguish generic thermodynamic mechanisms from system-specific molecular details and provide a foundation for understanding condensate regulation across diverse biological and synthetic systems.

Here, we address this question by developing a minimal Cahn–Hilliard–Flory–Huggins continuum model in which competing LLPS-promoting and LLPS-inhibiting interactions are encoded through a concentration-dependent interaction parameter. We show that this simple thermodynamic mechanism reproduces experimentally observed biphasic regulation in Tau– tannic acid and TDP-43–bis-ANS systems, captures morphology transitions and coarsening dynamics, and demonstrates that re-entrant condensate formation and dissolution emerge naturally from a concentration-dependent free-energy landscape.

## Model and Methods

To identify the minimal thermodynamic ingredients underlying re-entrant LLPS, we develop a continuum description within a Cahn–Hilliard–Flory–Huggins framework. The system is represented as a binary mixture of proteins and solvent, where the conserved order parameter *ϕ*(**r**, *t*) denotes the local protein volume fraction. The free energy combines the entropic contribution associated with mixing, effective protein–protein interactions encoded through a Flory interaction parameter, and an interfacial penalty accounting for concentration gradients. The corresponding free-energy functional is

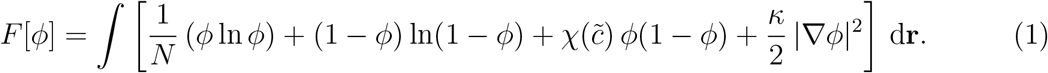

where *N* denotes the effective polymerization index of the protein, *κ* controls the interfacial energy, and 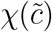 is an effective interaction parameter that encodes the competing LLPS-promoting and LLPS-inhibiting effects of small molecules.

To capture re-entrant LLPS, we introduce a concentration-dependent Flory interaction parameter,

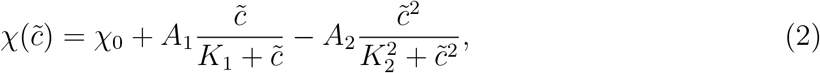

where *χ*_0_ is the intrinsic protein–solvent interaction parameter and 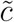 denotes the local effective small-molecule concentration. The first term represents a saturating LLPS-promoting contribution dominant at low concentrations, while the second term captures inhibitory effects arising from interaction screening or preferential solvation at higher concentrations. The parameters *A*_1_ and *A*_2_ determine the strengths of the competing interactions, whereas *K*_1_ and *K*_2_ set the corresponding crossover concentrations.

The LLPS-promoting contribution phenomenologically accounts for the enhancement of effective protein–protein attraction by small molecules at low concentrations and saturates as available interaction sites become occupied. In contrast, the inhibitory contribution captures the suppression of phase separation at higher concentrations through mechanisms such as preferential solvation and interaction screening. Their competition generates a nonmonotonic dependence of *χ* on modulator concentration, with the effective interaction strength becoming maximal only within a finite concentration window. As a result, the model naturally reproduces the experimentally observed enhancement of condensate formation at low concentrations and its suppression at higher concentrations. The physical interpretation of these competing effects is summarized schematically in Figure 1.

**Figure 1.**
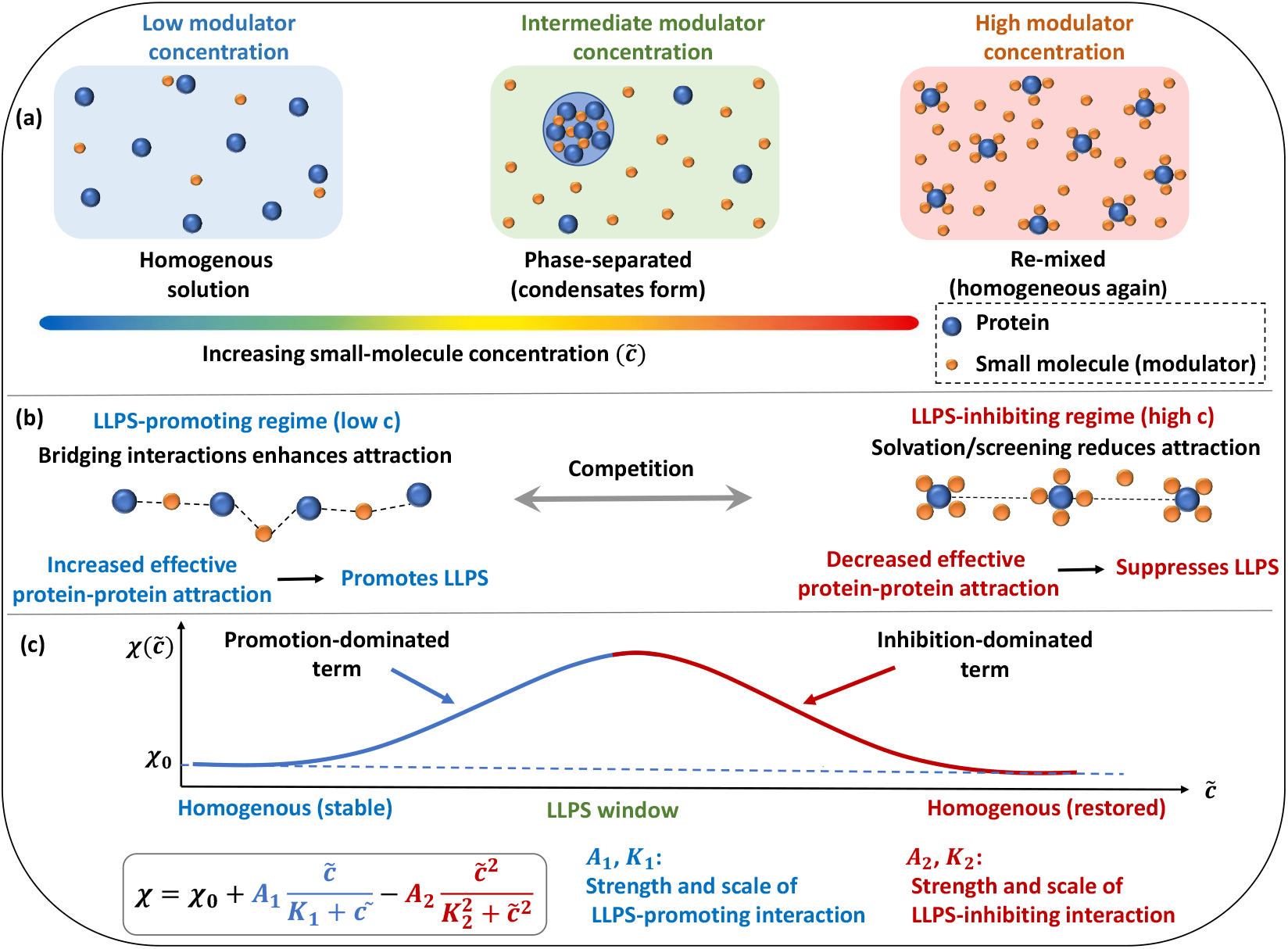
Schematic representation of re-entrant regulation of liquid–liquid phase separation by small molecules. (a) Increasing modulator concentration produces a biphasic response, with condensate formation at intermediate concentrations and remixing at high concentrations. (b) Illustration of the competing effects of small molecules on protein interactions: promotion of condensation at low concentrations and suppression of condensation at high concentrations. (c) Nonmonotonic dependence of the effective interaction parameter 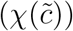 on modulator concentration arising from competing LLPS-promoting and LLPS-inhibiting contributions. The resulting concentration window between 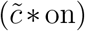 and 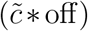 corresponds to conditions under which phase separation is observed.

Importantly, the proposed mechanism does not rely on system-specific molecular details. Instead, it assumes only that small molecules exert competing effects on intermolecular interactions, enhancing condensation at low concentrations and suppressing it at higher concentrations. The resulting nonmonotonic interaction landscape provides a generic route to re-entrant LLPS that can be applied across diverse protein–small-molecule systems.

The concentration-dependent interaction parameter enters the dynamics through the local chemical potential,

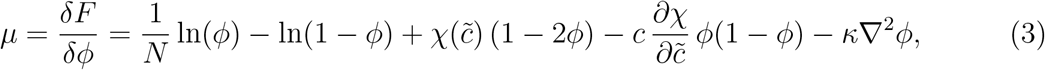

where

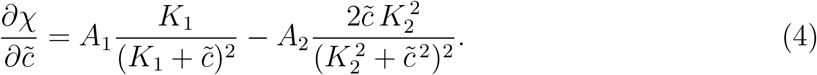

The additional term proportional to 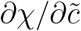 originates from the composition dependence of the interaction parameter and captures the feedback between local condensate composition and small-molecule regulation.

The order parameter evolves according to the Cahn-Hilliard equation,

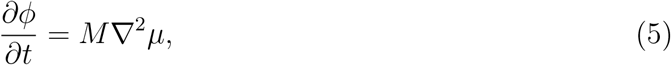

where *M* is the mobility. To account for preferential partitioning of small molecules into the dilute phase, we assume a local coupling of the form 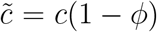, where *c* denotes the mean small-molecule concentration. This phenomenological ansatz reflects the tendency of many modulators to remain enriched in solvent-rich regions rather than partition uniformly into dense condensates. Consequently, the effective concentration available to regulate inter-molecular interactions decreases with increasing local protein density. The coupling therefore introduces a simple feedback between condensate formation and small-molecule regulation while avoiding the need to explicitly evolve a second conserved concentration field.

Numerical simulations were performed in two dimensions on a square domain with periodic boundary conditions using the conserved Cahn–Hilliard dynamics described by Eq. (5). Spatial derivatives were discretized using a second-order central finite-difference scheme, and the resulting evolution equations were integrated in time using an explicit Euler method. Unless otherwise stated, simulations were carried out on a domain of size *L*_*x*_ = *L*_*y*_ = 150 with spatial discretization Δ*x* = Δ*y* = 0.5 and time step Δ*t* = 10^*−*4^. The order parameter field was initialized with small-amplitude random fluctuations about a prescribed mean protein concentration 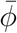, which remained conserved throughout the evolution. Unless otherwise specified, the mobility was fixed at *M* = 1.0. Simulations were evolved until steady-state morphologies were obtained or until sufficient temporal information was acquired for the analysis of phase-separation kinetics.

## Result and discussion

### Re-entrant LLPS in the Tau–Tannic acid system

We first consider the Tau–tannic acid system, which has recently emerged as a representative example of biphasic condensate regulation by small molecules. ^33^ Experiments have shown that tannic acid promotes Tau condensation at low concentrations while suppressing LLPS at higher concentrations, giving rise to a characteristic re-entrant phase diagram. Our objective is not to reproduce the microscopic chemistry of tannic acid binding in detail, but rather to determine whether the experimentally observed phase behavior can be explained using the minimal thermodynamic framework developed above. To this end, the parameters entering Eq. (2) were chosen such that the concentration dependence of the effective interaction parameter qualitatively reflects the experimentally observed turbidity trends. Unless stated otherwise, simulations were performed using *χ*_0_ = 0.19, *A*_1_ = 5.0, *K*_1_ = 1.0, *A*_2_ = 8.0, *K*_2_ = 3.7, *κ* = 10.0, and *M* = 1.0.

Representative concentration fields in Figure 2(a) illustrate the evolution of condensate morphology with increasing tannic acid concentration. In the absence of modulators, the system remains spatially homogeneous and does not undergo spontaneous demixing. Upon increasing *c*, concentration fluctuations become amplified and eventually develop into coexisting protein-rich and protein-poor domains. These domains subsequently coarsen and acquire increasingly sharp interfaces, indicating the emergence of a thermodynamically stable phase-separated state. The strongest compositional segregation is observed at intermediate concentrations, where well-defined condensates occupy a substantial fraction of the system. At larger values of *c*, however, the phase contrast progressively diminishes and the system ultimately returns to a homogeneous mixed state.

**Figure 2.**
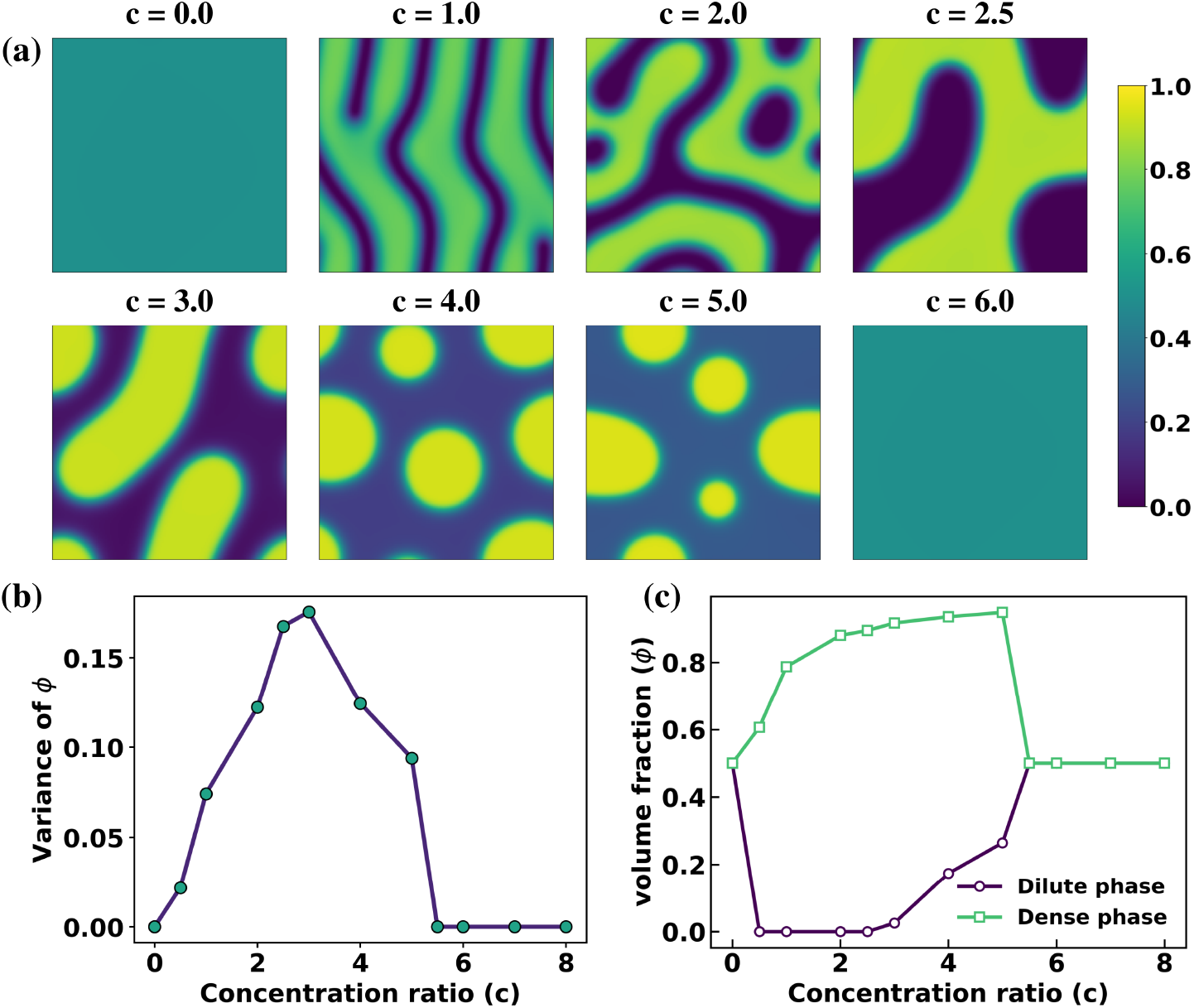
Biphasic regulation of liquid-liquid phase separation in the Tau–tannic acid system at equimolar composition 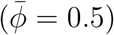. (a) Representative concentration fields *ϕ*(*x, y*) for increasing tannic acid to Tau concentration ratio *c*, showing the emergence of phase separation at intermediate *c* and its suppression at both low and high concentrations at *t* = 500. (b) Variance of the concentration field as a function of *c*, quantifying the nonmonotonic, re-entrant phase behavior. (c) Dilute- and dense-phase volume fractions as functions of *c*, indicating that the extent of phase separation, quantified by the difference between the coexisting phases, is maximal at intermediate concentrations.

This nonmonotonic evolution arises directly from the concentration dependence of the effective interaction parameter 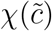. At low concentrations, the LLPS-promoting contribution dominates, strengthening effective protein–protein attractions and driving phase separation. As the modulator concentration increases, the inhibitory contribution becomes increasingly important, progressively reducing the thermodynamic driving force for demixing. Beyond a critical concentration, coexistence between protein-rich and protein-poor phases is no longer thermodynamically favorable, resulting in complete remixing of the system. The simulations therefore reproduce the experimentally observed biphasic response through a simple competition between LLPS-promoting and LLPS-inhibiting interactions, yielding a finite concentration window in which condensate formation is favored.

To quantify the extent of phase separation, we compute the variance of the order parameter,

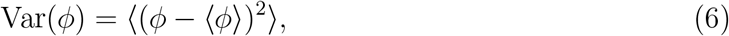

which serves as a computational analogue of experimental turbidity. Large values of Var(*ϕ*) correspond to strong spatial heterogeneity and pronounced coexistence between protein-rich and protein-poor regions, whereas values approaching zero indicate a homogeneous mixed state. As shown in Figure 2(b), the variance exhibits a pronounced nonmonotonic dependence on the tannic acid concentration, increasing upon entry into the phase-separated regime, reaching a maximum at intermediate concentrations, and subsequently decreasing as the system remixes. This behavior quantitatively confirms the biphasic response inferred from the morphology evolution.

Additional insight is provided by the coexistence compositions of the dilute and dense phases shown in Figure 2(c). The phase compositions separate progressively with increasing concentration, attain their largest contrast within the strongly phase-separated regime, and converge again at higher concentrations as the system approaches remixing. The non-monotonic behavior of both the variance and coexistence compositions demonstrates that condensate stability and phase contrast are maximized only within a finite concentration window, consistent with the experimentally observed re-entrant regulation of Tau condensation.

### Spinodal stability analysis and re-entrant phase boundaries

To establish the thermodynamic origin of the re-entrant behavior, we perform a spinodal stability analysis of the free-energy functional. Because the coupling 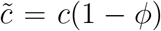 renders the interaction parameter *χ* composition dependent, additional coupling contributions enter the stability criterion,

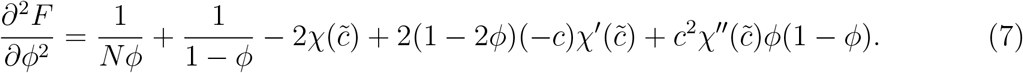

The spinodal condition, *∂*^2^*F/∂ϕ*^2^ = 0, defines the boundary between thermodynamically stable and unstable states in the (*ϕ, c*) plane. Whereas the numerical simulations reveal the morphology that emerges following phase separation, the spinodal analysis identifies the regions in parameter space where concentration fluctuations become intrinsically unstable.

The resulting phase diagram is shown in Figure 3(a). The spinodal boundary forms a closed instability region, indicating that LLPS is thermodynamically favorable only over a finite range of modulator concentrations. For the equimolar composition 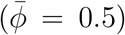, the homogeneous state first becomes unstable near *c* ≈ 0.35, while stability is recovered at *c* ≈ 5.44, leading to re-entrant mixing at larger concentrations. Figure 3(b) further demonstrates that *∂*^2^*F/∂ϕ*^2^ becomes negative only within this interval, precisely corresponding to the concentration range over which enhanced phase separation is observed in the simulations.

**Figure 3.**
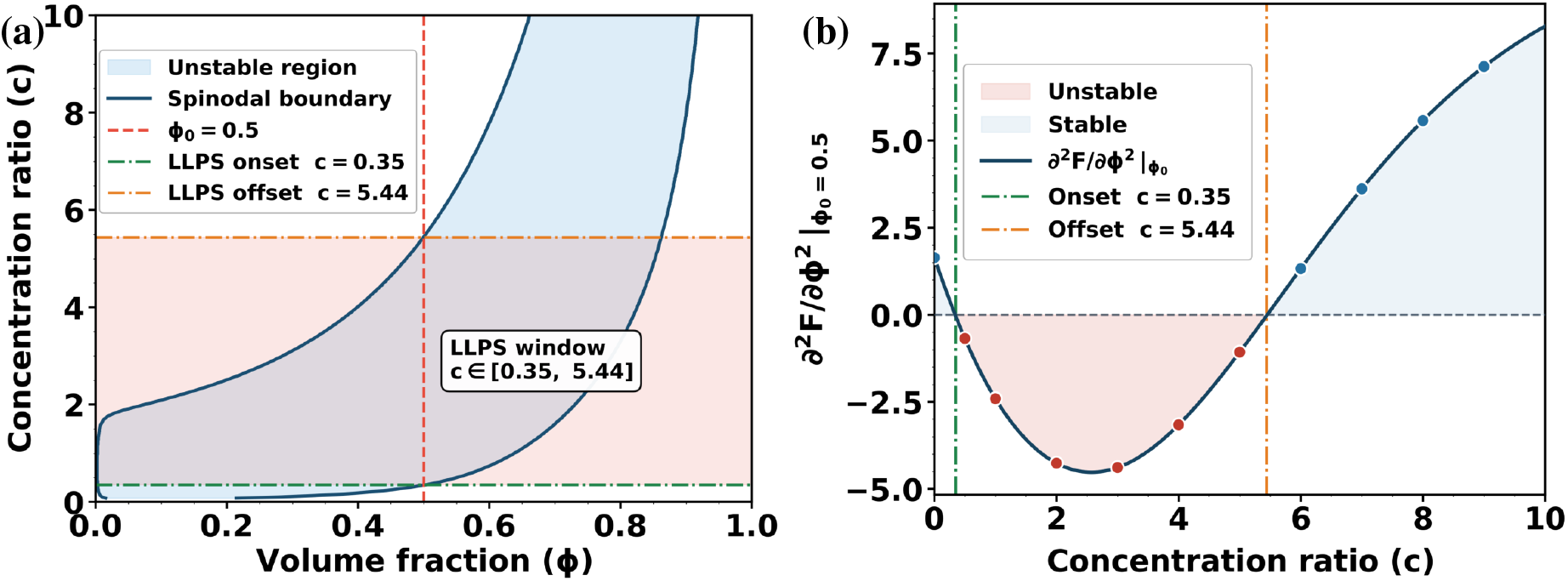
Phase stability analysis for the Tau–tannic acid system at 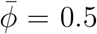. (a) Spinodal boundary in the (*ϕ, c*) plane; the shaded blue region marks the thermodynamically unstable regime (*∂*^2^*F/∂ϕ*^2^ *<* 0). Intersections of the 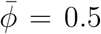 line (red dashed) with the boundary define the LLPS onset (*c* ≈ 0.35) and offset (*c* ≈ 5.44). (b) *∂*^2^*F/∂ϕ*^2^ evaluated at 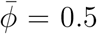 versus *c*. The quantity is negative (red-shaded) within the predicted window, confirming spinodal instability and re-entrant mixing at high *c*. Filled circles mark representative *c* values used in simulations.

The lower spinodal boundary marks the onset of instability driven by the LLPS-promoting contribution to 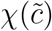, whereas the upper boundary reflects the increasing influence of LLPS-inhibiting interactions that ultimately restore thermodynamic stability. These results demonstrate that the re-entrant transition is not merely a kinetic phenomenon but arises directly from the underlying free-energy landscape encoded in the concentration-dependent interaction parameter.

### Condensate formation in the dilute regime

We next examine a more dilute system with mean protein concentration 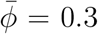, which is closer to the conditions under which many biomolecular condensates form in vivo. In cellular environments, condensates often occupy only a small fraction of the available volume and coexist with a large solvent-rich background. It is therefore important to determine whether the re-entrant mechanism identified above remains operative away from the equimolar regime.

Representative concentration fields are shown in Figure 4(a). In contrast to the morphologies observed at 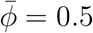, phase separation now produces discrete protein-rich droplets dispersed within a continuous dilute phase. Despite this change in morphology, the overall response to tannic acid concentration remains qualitatively unchanged. At low concentrations, the system remains homogeneous, while increasing *c* promotes droplet nucleation and growth. The most pronounced phase separation occurs at intermediate concentrations, where well-defined condensates coexist with a dilute background. Upon further increasing *c*, the droplets progressively dissolve and the system ultimately returns to a homogeneous mixed state.

**Figure 4.**
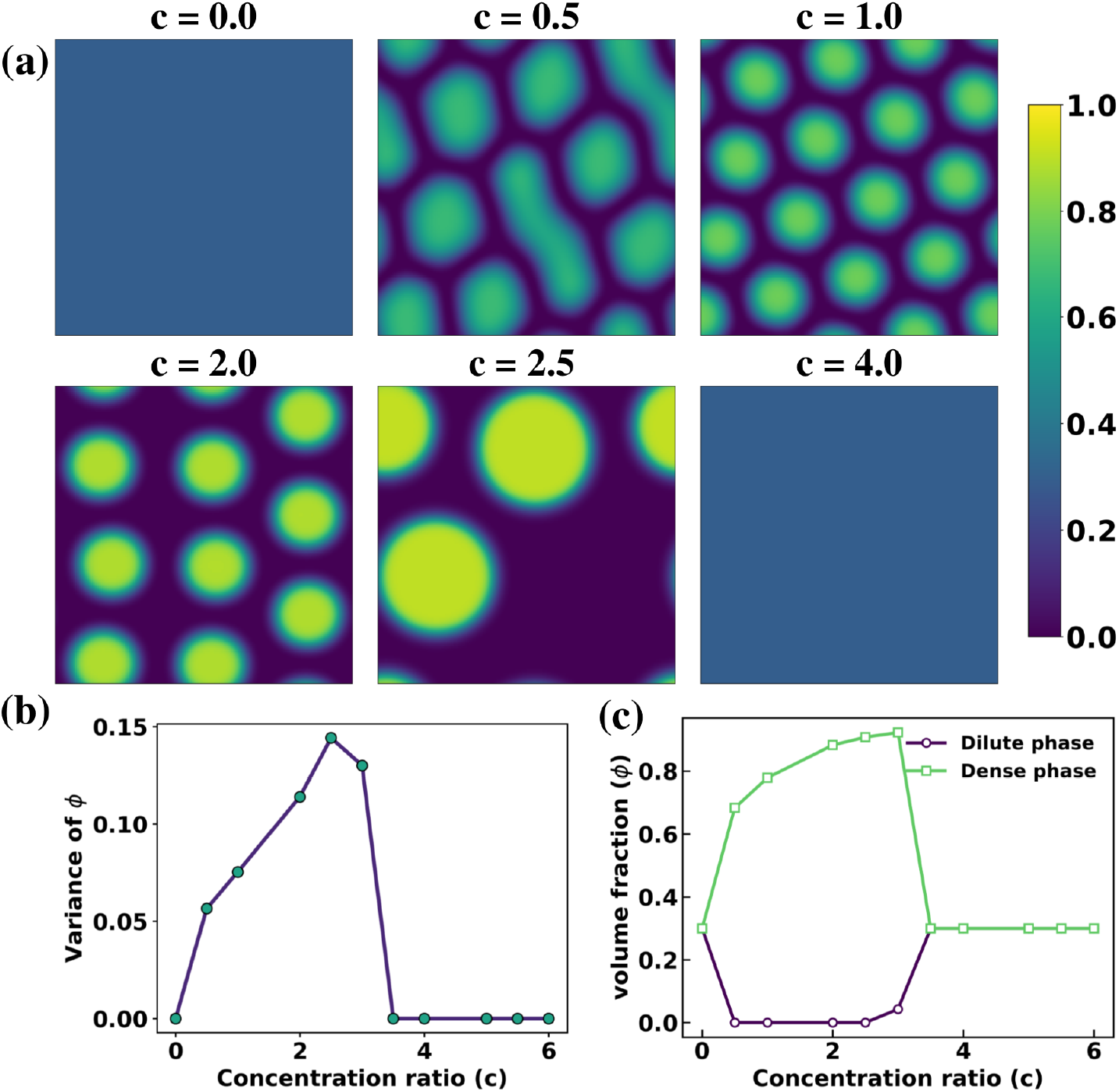
Biphasic modulation of liquid-liquid phase separation in the Tau–tannic acid system under dilute conditions 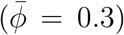. (a) Concentration fields *ϕ*(*x, y*) for increasing tannic acid to Tau concentration ratio *c* at *t* = 500. (b) Variance of *ϕ* as a function of *c*, showing a nonmonotonic response. (c) Coexisting dilute- and dense-phase volume fractions, with the largest phase contrast occurring at intermediate *c*.

The corresponding variance and coexistence compositions are shown in Figures 4(b,c). Both quantities exhibit the same nonmonotonic dependence on modulator concentration observed in the equimolar regime. The variance reaches a maximum within the phase-separated region and subsequently decreases as the system undergoes re-entrant mixing. Likewise, the compositional contrast between the dense and dilute phases is greatest at intermediate concentrations and vanishes outside the coexistence regime. These results demonstrate that the biphasic regulation mechanism is robust with respect to the overall protein concentration and governs both interconnected domain morphologies and isolated droplet states.

The corresponding spinodal phase diagram for the dilute regime is provided in the Supporting Information (Figure S1), where the instability region again forms a closed domain in the (*ϕ, c*) plane. Together, these results indicate that the finite concentration window for LLPS is a generic consequence of the concentration-dependent interaction parameter and the resulting free-energy landscape, rather than a feature specific to a particular average composition.

### Domain growth and coarsening dynamics

To further characterize the kinetics of phase separation, we quantify the characteristic domain size *R*(*t*) from the structure factor 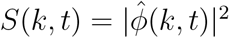, where 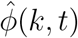 is the Fourier transform of the fluctuation field *δϕ* = *ϕ* − ⟨*ϕ*⟩. The characteristic length scale is defined as

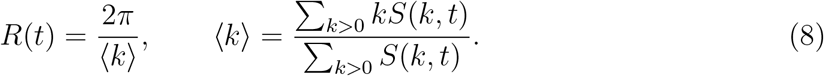

While the spinodal analysis identifies the concentration range over which phase separation is thermodynamically favorable, the temporal evolution of *R*(*t*) provides insight into the subsequent growth and maturation of condensates. Figure 5 shows the evolution of the characteristic domain size for representative modulator concentrations. At low and high concentrations, corresponding to thermodynamically stable homogeneous states, *R*(*t*) remains nearly constant throughout the simulation. In contrast, intermediate concentrations exhibit sustained domain growth arising from the coarsening of protein-rich condensates through diffusive transport and interfacial energy minimization. These results demonstrate that the same concentration window responsible for thermodynamic instability also governs the efficiency of mesoscale condensate growth.

**Figure 5.**
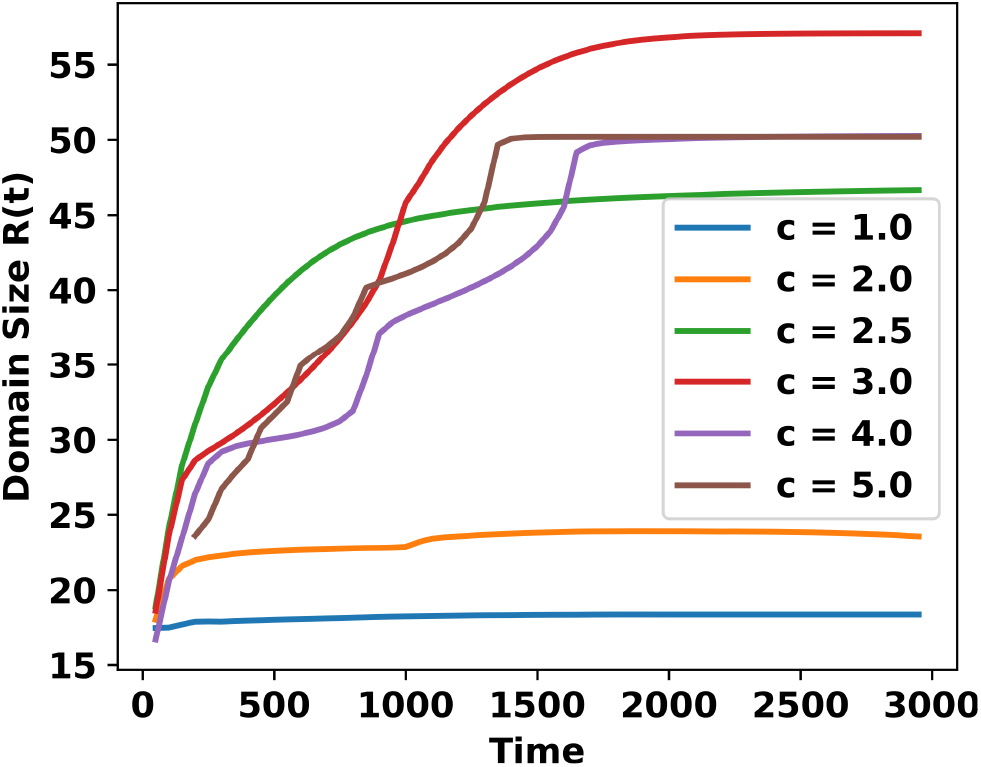
Temporal evolution of the characteristic domain size *R*(*t*) for the Tau–tannic acid system at 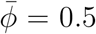 for representative small-molecule concentrations *c*. At intermediate concentrations, *R*(*t*) exhibits power-law growth consistent with diffusive coarsening, while low and high concentrations suppress domain growth due to the absence of stable phase separation.

At late times, the domain size follows an approximate power-law growth, *R*(*t*) ∼ *t*^*n*^, with the extracted exponent approaching the Lifshitz–Slyozov–Wagner prediction *n* ≈ 1*/*3 for diffusion-limited coarsening in conserved systems. ^38,39^ The agreement with classical coarsening theory indicates that, although small molecules strongly regulate the onset and extent of phase separation, the subsequent evolution of condensates remains governed by universal diffusive transport mechanisms. Thus, re-entrant modulation primarily determines when and over what concentration range condensates can form, while the late-stage growth kinetics retain the characteristic behavior of conserved order-parameter systems. More broadly, these results highlight that small-molecule regulation influences not only the thermodynamic stability of biomolecular condensates but also the temporal window over which condensate growth and maturation can proceed.

### Transferability of the framework: The bis-ANS–TDP-43 system

To examine whether the proposed mechanism is specific to the Tau–tannic acid system or represents a more general principle of chemically regulated condensate formation, we next apply the same continuum framework to the bis-ANS–TDP-43 system. Experimental studies have shown that bis-ANS regulates TDP-43 phase separation in a biphasic manner, promoting condensate formation at low concentrations while suppressing LLPS at higher concentrations.^31^ This system therefore provides an independent test of whether re-entrant phase behavior can emerge from the competition between LLPS-promoting and LLPS-inhibiting interactions encoded in Eq. (2).

Without modifying the governing equations, system-specific behavior was incorporated solely through the parameters entering the concentration-dependent interaction parameter 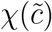. Guided by the experimentally observed turbidity profiles, we employed *χ*_0_ = 0.19, *A*_1_ = 5.0, *K*_1_ = 1.25, *A*_2_ = 6.2, *K*_2_ = 2.7, *κ* = 8.0, and *M* = 1.0. Representative concentration fields together with the corresponding variance and coexistence curves are shown in Figure 6.

**Figure 6.**
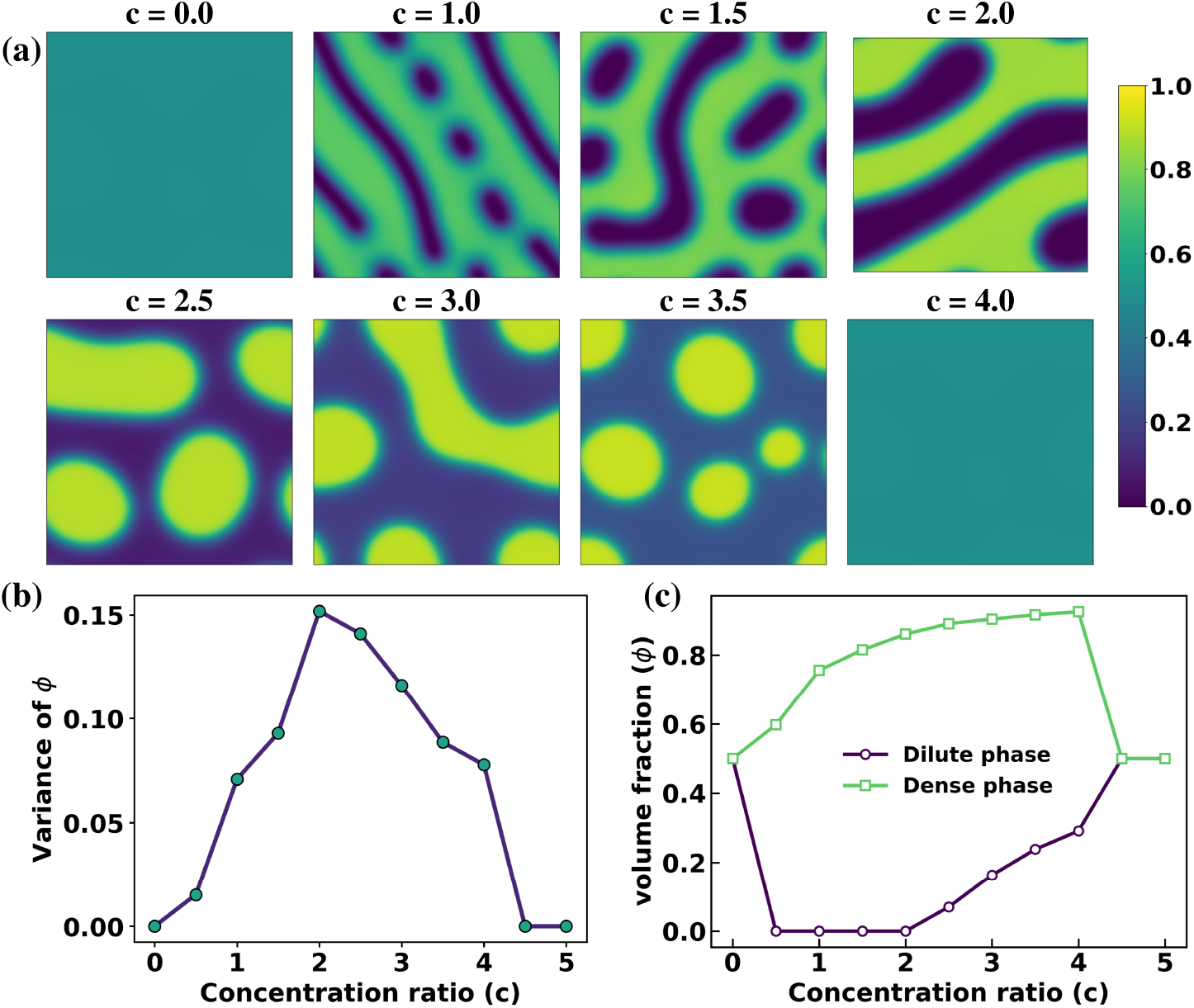
Biphasic LLPS in the bis-ANS–TDP-43 system at 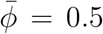. (a) Concentration fields *ϕ*(*x, y*) at *t* = 500 for increasing *c*. (b) Variance of *ϕ* versus *c*, exhibiting a pronounced maximum near *c* ≈ 2.5. (c) Coexisting dilute- and dense-phase volume fractions versus *c*.

The simulations reproduce the characteristic biphasic response observed experimentally. At low bis-ANS concentrations, the system remains nearly homogeneous. Increasing the modulator concentration induces phase separation and the emergence of protein-rich condensates, with the strongest phase contrast occurring at intermediate concentrations. Upon further increasing *c*, the inhibitory contribution to 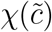 becomes dominant, resulting in condensate dissolution and restoration of a homogeneous mixed state. Consistent with these morphological observations, the variance of the order parameter exhibits a pronounced maximum and the coexistence compositions display the largest contrast within the phase-separated regime.

To establish the thermodynamic origin of this behavior, we additionally performed a spinodal stability analysis using the same criterion as for the Tau–tannic acid system. The resulting phase diagram for the equimolar composition (Figure S2) exhibits a closed instability region within which *∂*^2^*F/∂ϕ*^2^ *<* 0, indicating spontaneous amplification of concentration fluctuations. The boundaries of this instability window closely coincide with the concentration range over which the variance is enhanced, confirming that the observed biphasic response originates directly from the underlying free-energy landscape. Thus, despite the distinct molecular nature of the modulator, the same thermodynamic mechanism governs the emergence and disappearance of phase separation.

To further assess the robustness of the predicted re-entrant behavior, we examined a dilute regime representative of condensates embedded within a solvent-rich environment. Representative concentration fields, variance profiles, and coexistence compositions are shown in Figure S3, while the corresponding spinodal phase diagram is presented in Figure S4. Similar to the Tau–tannic acid system, the model predicts a finite concentration window for LLPS together with re-entrant mixing at larger modulator concentrations. The persistence of this behavior across both equimolar and dilute compositions demonstrates that the essential features of the biphasic response are governed primarily by the concentration dependence of the effective interaction parameter rather than the average protein concentration.

Importantly, no modification of the continuum equations was required when moving from Tau–tannic acid to bis-ANS–TDP-43. The same theoretical framework therefore captures re-entrant condensate formation across chemically distinct protein–small-molecule systems using only a small set of physically interpretable parameters. This transferability suggests that competition between LLPS-promoting and LLPS-inhibiting interactions may constitute a generic thermodynamic mechanism underlying chemically regulated biomolecular condensates.

### Robustness with respect to mobility

To assess the extent to which the predicted biphasic response depends on transport kinetics, we examined both constant and composition-dependent mobilities within the same thermodynamic framework. While the mobility influences the rate of domain growth and the detailed morphology evolution, it does not alter the underlying free-energy landscape responsible for phase separation. Consequently, comparing different mobility functions provides a useful means of distinguishing thermodynamic effects from kinetic ones.

In addition to the constant mobility used throughout the preceding sections (*M* = 1.0), we considered a composition-dependent mobility of the form

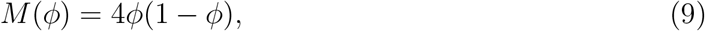

which vanishes in the pure phases and attains its maximum at intermediate compositions. Such interfacial mobilities are widely employed in phase-field models because they suppress diffusion within bulk phases while enhancing transport near interfaces.^40–42^

Figures 7 compare the resulting phase behavior for the Tau–tannic acid system. In both cases, the morphology evolution, variance profiles, and coexistence compositions exhibit the same characteristic biphasic dependence on modulator concentration. The variance of the order parameter displays a pronounced maximum at intermediate concentrations, while phase separation disappears at both low and high concentrations, consistent with the reentrant behavior observed throughout this study. The primary effect of the composition-dependent mobility is kinetic rather than thermodynamic. Relative to the constant-mobility case, domain growth proceeds more slowly and the system remains less coarsened at comparable simulation times, leading to reduced compositional contrast between the coexisting phases. Nevertheless, the concentration window over which phase separation occurs remains essentially unchanged. These results demonstrate that the emergence of re-entrant LLPS is governed primarily by the free-energy landscape encoded in the concentration-dependent interaction parameter 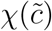 and is robust with respect to the specific choice of mobility.

**Figure 7.**
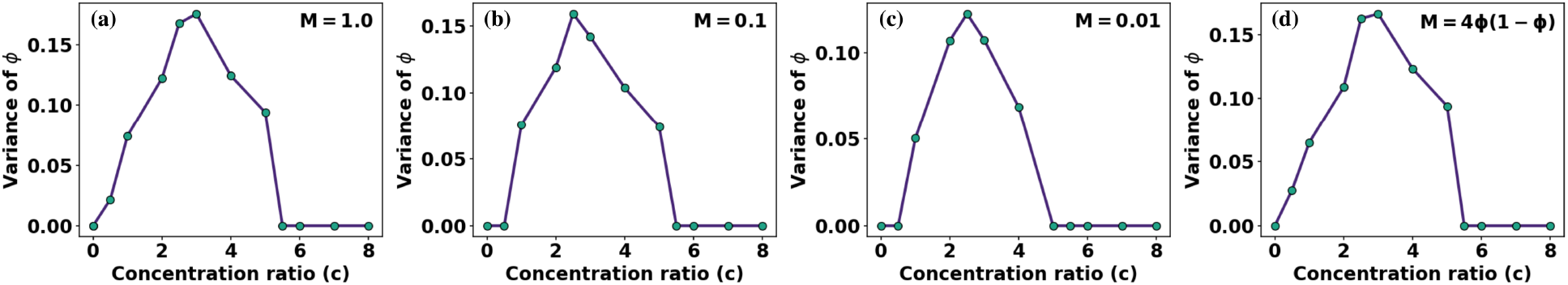
Effect of mobility on biphasic LLPS in the Tau–tannic acid system. Variance of the order parameter, Var(*ϕ*), as a function of small-molecule concentration *c* for (a) *M* = 1.0, (b) *M* = 0.10, (c) *M* = 0.01, and (d) *M* (*ϕ*) = 4*ϕ*(1 − *ϕ*). The persistence of the nonmonotonic maximum across all mobility choices confirms that the re-entrant behavior is thermodynamically driven and robust to variations in transport kinetics.

## Concluding remarks

In this work, we have developed a minimal continuum theory for the re-entrant regulation of liquid-liquid phase separation by small molecules within a Cahn-Hilliard-Flory-Huggins framework. The central element of the model is a concentration-dependent interaction parameter, 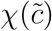, that captures the competition between LLPS-promoting and LLPS-inhibiting interactions. This simple thermodynamic mechanism generates a nonmonotonic dependence of condensate stability on modulator concentration and naturally gives rise to the experimentally observed biphasic response.

Application of the framework to the Tau–tannic acid and bis-ANS–TDP-43 systems demonstrates that a common mesoscale description can account for condensate formation and dissolution across chemically distinct protein–small-molecule combinations. The model reproduces finite concentration windows for phase separation, predicts the associated spinodal boundaries, and captures morphology evolution, coexistence behavior, and domain coarsening dynamics. The close correspondence between numerical simulations and thermodynamic stability analysis establishes that re-entrant mixing emerges directly from the underlying free-energy landscape rather than from kinetic effects.

A notable feature of the present approach is its transferability. Without modification of the governing continuum equations, the same framework successfully describes multiple experimentally studied systems using only a small set of physically interpretable parameters. Together with the robustness observed for different mobility functions, these results suggest that competition between condensate-promoting and condensate-inhibiting interactions constitutes a generic thermodynamic mechanism underlying chemically regulated biomolecular phase separation.

Beyond the specific systems considered here, the framework provides a versatile platform for investigating a broad range of problems involving condensate regulation. Future extensions may incorporate multicomponent mixtures, reaction-coupled nonequilibrium processes, chemically programmable phase separation, and active biomolecular assemblies. More generally, the present theory establishes a mesoscale link between molecular-scale regulation and emergent condensate behavior, offering a physically transparent framework for understanding how small molecules modulate biomolecular phase separation and for guiding future strategies aimed at controlling pathological condensate formation associated with neurodegenerative disease and related disorders.

## Author Contributions

Both the authors contributed equally to the conceptualization of the study, development of the theoretical framework, numerical simulations, analysis of the results, and writing of the manuscript. Both authors discussed the results and approved the final version of the manuscript.

## Data availability

The data supporting this article have been included as part of the Supporting Information (SI). SI provides additional results complementing the main text. Section S1 presents the spinodal phase diagram for the Tau–tannic acid system at 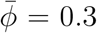. Section S2 applies the same framework to the bis-ANS–TDP-43 system for dilute conditions, including spinodal analysis. SI contains Supplementary Figures S1–S4 and Section S3 provides corresponding video files. The simulation code used in this work is publicly available at GitHub: https://github.com/pushpita-group/LLPS.git.

## Acknowledgement

The authors acknowledge IISER Thiruvananthapuram (IISERTVM) for the computational resources. Achal Jadhav thanks IISERTVM for the Institute fellowship.

## References

(1) Dolgin, E. What lava lamps and vinaigrette can teach us about cell biology. Nature 2018, 555, 300–303.

(2) Brangwynne, C. P.; Eckmann, C. R.; Courson, D. S.; Rybarska, A.; Hoege, C.; Gharakhani, J.; Jülicher, F.; Hyman, A. A. Germline P granules are liquid droplets that localize by controlled dissolution/condensation. Science 2009, 324, 1729–1732.

(3) Brangwynne, C. P.; Mitchison, T. J.; Hyman, A. A. Active liquid-like behavior of nucleoli determines their size and shape in Xenopus laevis oocytes. Proceedings of the National Academy of Sciences 2011, 108, 4334–4339.

(4) Nott, T. J.; Petsalaki, E.; Farber, P.; Jervis, D.; Fussner, E.; Plochowietz, A.; Craggs, T. D.; Bazett-Jones, D. P.; Pawson, T.; Forman-Kay, J. D., et al. Phase transition of a disordered nuage protein generates environmentally responsive membraneless organelles. Molecular cell 2015, 57, 936–947.

(5) Hyman, A. A.; Weber, C. A.; Jülicher, F. Liquid-liquid phase separation in biology. Annual review of cell and developmental biology 2014, 30, 39–58.

(6) Berry, J.; Brangwynne, C. P.; Haataja, M. Physical principles of intracellular organization via active and passive phase transitions. Reports on Progress in Physics 2018, 81, 046601.

(7) Weber, C. A.; Zwicker, D.; Jülicher, F.; Lee, C. F. Physics of active emulsions. Reports on Progress in Physics 2019, 82, 064601.

(8) Jülicher, F.; Weber, C. A. Droplet physics and intracellular phase separation. Annual Review of Condensed Matter Physics 2024, 15, 237–261.

(9) Zwicker, D.; Paulin, O. W.; Ter Burg, C. Physics of droplet regulation in biological cells. Reports on Progress in Physics 2025, 88, 116601.

(10) Saar, K. L.; Qian, D.; Good, L. L.; Morgunov, A. S.; Collepardo-Guevara, R.; Best, R. B.; Knowles, T. P. Theoretical and data-driven approaches for biomolecular condensates. Chemical Reviews 2023, 123, 8988–9009.

(11) Zhou, H.-X.; Kota, D.; Qin, S.; Prasad, R. Fundamental aspects of phase-separated biomolecular condensates. Chemical Reviews 2024, 124, 8550–8595.

(12) Buchan, J. R.; Parker, R. Eukaryotic stress granules: the ins and outs of translation. Molecular cell 2009, 36, 932–941.

(13) Decker, C. J.; Parker, R. P-bodies and stress granules: possible roles in the control of translation and mRNA degradation. Cold Spring Harbor perspectives in biology 2012, 4, a012286.

(14) Boisvert, F.-M.; Van Koningsbruggen, S.; Navascués, J.; Lamond, A. I. The multifunctional nucleolus. Nature reviews Molecular cell biology 2007, 8, 574–585.

(15) Gall, J. G. The centennial of the Cajal body. Nature reviews Molecular cell biology 2003, 4, 975–980.

(16) Prasad, A.; Bharathi, V.; Sivalingam, V.; Girdhar, A.; Patel, B. K. Molecular mechanisms of TDP-43 misfolding and pathology in amyotrophic lateral sclerosis. Frontiers in molecular neuroscience 2019, 12, 25.

(17) Li, P.; Chen, J.; Wang, X.; Su, Z.; Gao, M.; Huang, Y. Liquidliquid phase separation of tau: Driving forces, regulation, and biological implications. Neurobiology of Disease 2023, 183, 106167.

(18) Kilgore, H. R.; Young, R. A. Learning the chemical grammar of biomolecular condensates. Nature Chemical Biology 2022, 18, 1298–1306.

(19) Maruri-Lopez, I.; Chodasiewicz, M. Involvement of small molecules and metabolites in regulation of biomolecular condensate properties. Current Opinion in Plant Biology 2023, 74, 102385.

(20) Matsuzawa, T., et al. Metabolites Shift Equilibria of Biomolecular Condensates. bioRxiv 2026, 2026–01, Preprint.

(21) Kitamura, K.; Oshima, A.; Sasaki, F.; Shiramasa, Y.; Yamamoto, R.; Kameda, T.; Kitazawa, S.; Kitahara, R. Modulation of biomolecular liquid–liquid phase separation by preferential hydration and interaction of small osmolytes with proteins. The Journal of Physical Chemistry Letters 2024, 15, 7620–7627.

(22) Duan, C.; Wang, R. A unified description of salt effects on the liquid–liquid phase separation of proteins. ACS Central Science 2024, 10, 460–468.

(23) Kota, D.; Prasad, R.; Zhou, H.-X. Adenosine triphosphate mediates phase separation of disordered basic proteins by bridging intermolecular interaction networks. Journal of the American Chemical Society 2024, 146, 1326–1336.

(24) Lin, Y.-H., et al. Electrostatics of salt-dependent reentrant phase behaviors highlights diverse roles of ATP in biomolecular condensates. eLife 2025, 13, RP100284.

(25) Sarkar, S.; Mondal, J. Mechanistic Insights on ATP’s Role as a Hydrotrope. The Journal of Physical Chemistry B 2021, 125, 7717–7731.

(26) Patel, A.; Malinovska, L.; Saha, S.; Wang, J.; Alberti, S.; Krishnan, Y.; Hyman, A. A. ATP as a biological hydrotrope. Science 2017, 356, 753–756.

(27) Kang, J.; Lim, L.; Song, J. ATP enhances at low concentrations but dissolves at high concentrations liquid-liquid phase separation (LLPS) of ALS/FTD-causing FUS. Biochemical and biophysical research communications 2018, 504, 545–551.

(28) Polymenidou, M.; Lagier-Tourenne, C.; Hutt, K. R.; Huelga, S. C.; Moran, J.; Liang, T. Y.; Ling, S.-C.; Sun, E.; Wancewicz, E.; Mazur, C., et al. Long pre-mRNA depletion and RNA missplicing contribute to neuronal vulnerability from loss of TDP-43. Nature neuroscience 2011, 14, 459–468.

(29) Wang, Y.; Mandelkow, E. Tau in physiology and pathology. Nature reviews neuroscience 2016, 17, 22–35.

(30) Ballatore, C.; Lee, V. M.-Y.; Trojanowski, J. Q. Tau-mediated neurodegeneration in Alzheimer’s disease and related disorders. Nature reviews neuroscience 2007, 8, 663–672.

(31) Babinchak, W. M.; Dumm, B. K.; Venus, S.; Boyko, S.; Putnam, A. A.; Jankowsky, E.; Surewicz, W. K. Small molecules as potent biphasic modulators of protein liquid-liquid phase separation. Nature communications 2020, 11, 5574.

(32) Dang, M.; Lim, L.; Kang, J.; Song, J. ATP biphasically modulates LLPS of TDP-43 PLD by specifically binding arginine residues. Communications Biology 2021, 4, 714.

(33) Xiang, J.; Chen, J.; Liu, Y.; Ye, H.; Han, Y.; Li, P.; Gao, M.; Huang, Y. Tannic acid as a biphasic modulator of tau protein liquid–liquid phase separation. International Journal of Biological Macromolecules 2024, 275, 133578.

(34) Banerjee, P. R.; Milin, A. N.; Moosa, M. M.; Onuchic, P. L.; Deniz, A. A. Reentrant phase transition drives dynamic substructure formation in ribonucleoprotein droplets. Angewandte Chemie International Edition 2017, 56, 11354–11359.

(35) Milin, A. N.; Deniz, A. A. Reentrant phase transitions and non-equilibrium dynamics in membraneless organelles. Biochemistry 2018, 57, 2470–2477.

(36) Wurtz, J. D.; Lee, C. F. Stress granule formation via ATP depletion-triggered phase separation. New Journal of Physics 2018, 20, 045008.

(37) Ren, C.-L.; Shan, Y.; Zhang, P.; Ding, H.-M.; Ma, Y.-Q. Uncovering the molecular mechanism for dual effect of ATP on phase separation in FUS solution. Science Advances 2022, 8, eabo7885.

(38) Lifshitz, I.; Slyozov, V. The kinetics of precipitation from supersaturated solid solutions. Journal of Physics and Chemistry of Solids 1961, 19, 35–50.

(39) Wagner, C. Theorie der Alterung von Niederschlägen durch Umlösen (Ostwald-Reifung). Zeitschrift für Elektrochemie, Berichte der Bunsengesellschaft für physikalische Chemie 1961, 65, 581–591.

(40) Bray, A.; Emmott, C. Lifshitz-Slyozov scaling for late-stage coarsening with an order-parameter-dependent mobility. Physical Review B 1995, 52, R685.

(41) Dai, S.; Du, Q. Computational studies of coarsening rates for the Cahn–Hilliard equation with phase-dependent diffusion mobility. Journal of Computational Physics 2016, 310, 85–108.

(42) Dai, S.; Du, Q. Coarsening mechanism for systems governed by the Cahn–Hilliard equation with degenerate diffusion mobility. Multiscale Modeling & Simulation 2014, 12, 1870–1889.

