## Supplementary information for "A minimal thermodynamic theory for re-entrant liquid-liquid phase separation regulated by small molecules"

---

This Supplementary Information provides additional results complementing the main text. Section S1 presents the spinodal phase diagram for the Tau–tannic acid system at dilute regime $\bar{\phi} = 0.3$. Section S2 applies the same framework to the bis-ANS–TDP-43 system at both equimolar and dilute conditions, including spinodal analysis. Section S3 provides supplementary movies illustrating the time evolution of phase separation dynamics in the Tau–tannic acid system at both equimolar ($\bar{\phi} = 0.5$) and dilute ($\bar{\phi} = 0.3$) conditions.

### S1. Spinodal Analysis: Tau–Tannic Acid System at $\bar{\phi} = 0.3$

The spinodal boundary is determined by $\partial^2 f/\partial\phi^2 = 0$, where $f(\phi)$ is the Flory–Huggins bulk free energy density with concentration-dependent $\chi(\tilde{c})$. The region where $\partial^2 f/\partial\phi^2 < 0$ is linearly unstable and drives spontaneous phase separation.

As shown in Fig. S1, the system enters the unstable region at a lower onset concentration than the equimolar case, reflecting the shift in thermodynamic stability upon dilution. The spinodal window is in good qualitative agreement with the variance curve (Fig. 4, main text), confirming that the biphasic response is encoded in the free-energy landscape through the competition between the promoting and inhibitory contributions to $\chi(\tilde{c})$.

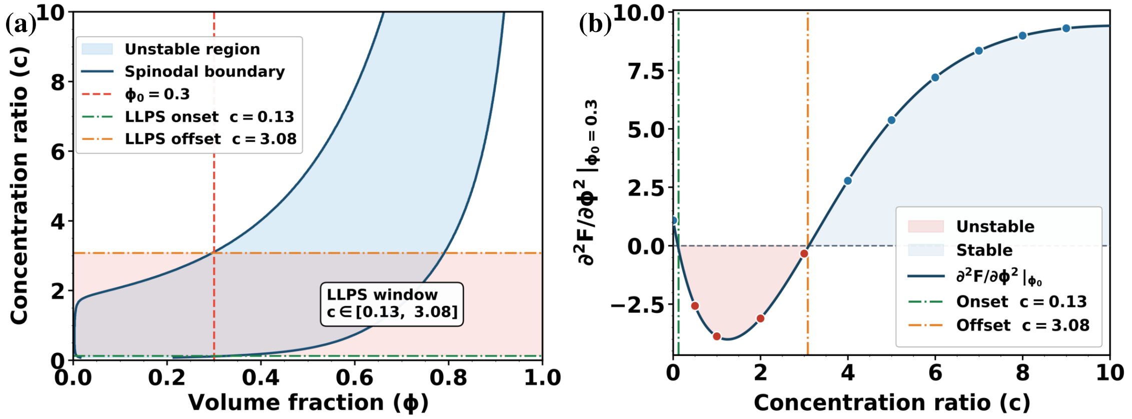


**Figure S1** Spinodal phase diagram for the Tau–tannic acid system at $\bar{\phi} = 0.3$. (a) Spinodal boundary in the $(\phi, c)$ plane; unstable region shaded; dashed line marks $\bar{\phi} = 0.3$. (b) $\partial^2 f/\partial\phi^2|_{\bar{\phi}=0.3}$ versus $c$, becoming negative within a finite window consistent with the variance peak in (Fig. 4, main text).

### S2. Biphasic LLPS Modulation in the bis-ANS–TDP-43 System

#### S2.1 Equimolar regime ($\bar{\phi} = 0.5$): spinodal analysis

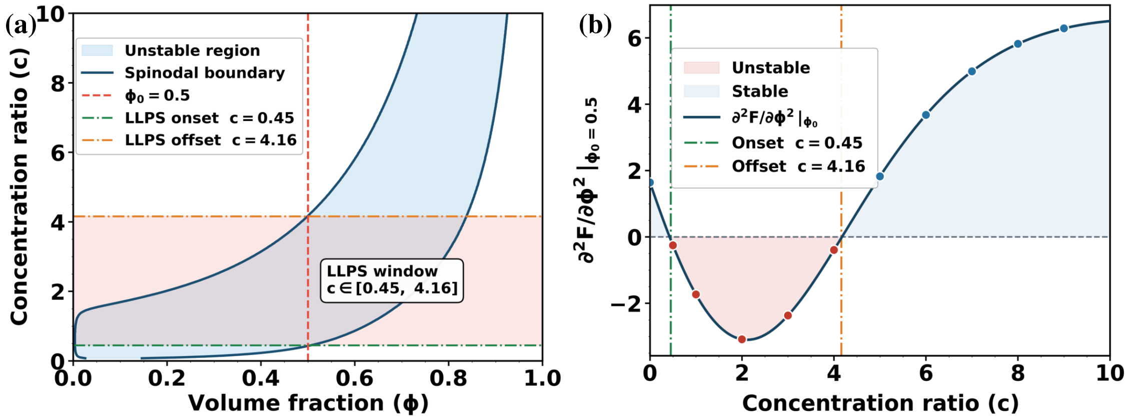


**Figure S2** Spinodal phase diagram for the bis-ANS–TDP-43 system at $\bar{\phi} = 0.5$. (a) Spinodal boundary in the $(\phi, c)$ plane; dashed line marks $\bar{\phi} = 0.5$. (b) $\partial^2 f/\partial\phi^2|_{\bar{\phi}=0.5}$ versus $c$, in agreement with the variance peak in (Fig. 6, main text).

As shown in Fig. S2, the spinodal window of instability is in good qualitative agreement with the variance peak (Fig. 6, main text), confirming that the biphasic numerical response is faithfully encoded in the thermodynamic free-energy landscape.

#### S2.2 Dilute regime ($\bar{\phi} = 0.3$): numerical simulations

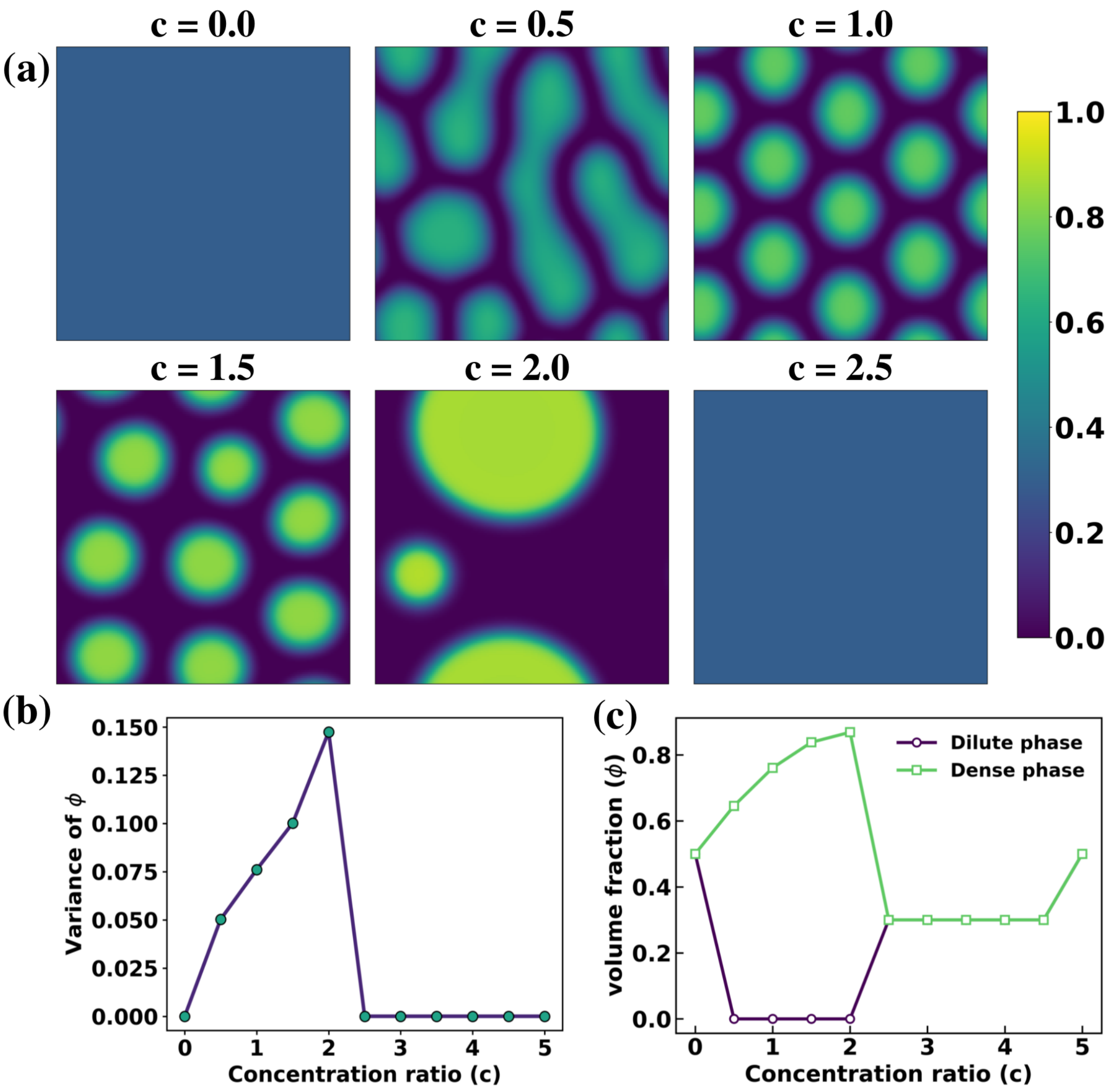


**Figure S3** Biphasic LLPS in the bis-ANS–TDP-43 system at $\bar{\phi} = 0.3$. (a) Concentration fields $\phi(x, y)$ at $t = 500$ for increasing $c$. (b) Variance of $\phi$ versus $c$, showing a nonmonotonic peak. (c) Coexisting dilute- and dense-phase volume fractions versus $c$.

Simulations at $\bar{\phi} = 0.3$ with $A_1 = 4.0$ and $A_2 = 4.0$ (all other parameters unchanged) reproduce the biphasic response (Fig. S3), with discrete protein-rich droplets forming at intermediate $c$ and re-entrant mixing at high $c$, consistent with the dilute Tau–tannic acid system (Fig. 4, main text).

#### S2.3. Dilute regime ($\bar{\phi} = 0.3$): spinodal analysis

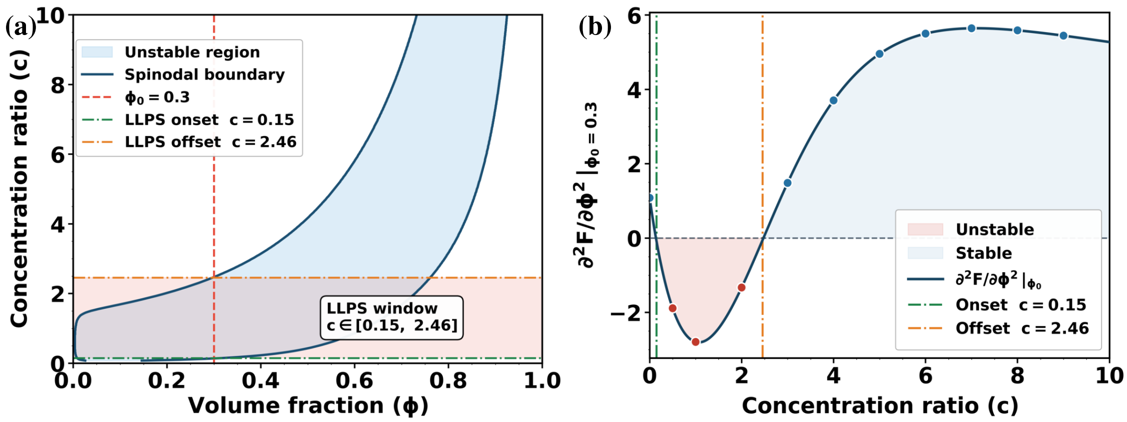


**Figure S4** Spinodal phase diagram for the bis-ANS–TDP-43 system at $\bar{\phi} = 0.3$. (a) Spinodal boundary in the $(\phi, c)$ plane; dashed line marks $\bar{\phi} = 0.3$. (b) $\partial^2 f/\partial\phi^2|_{\bar{\phi}=0.3}$ versus $c$, in agreement with the variance peak in Fig. S3(b).

Fig. S4 confirms a finite window of linear instability in good agreement with the variance (Fig. S3(b)). Together, Figs. S2–S4 establish that the free-energy landscape faithfully encodes the biphasic phase behavior across both concentration regimes and both protein–small-molecule systems.

### S3. Movies

All simulations were performed with $\chi_0 = 0.19$, $c = 2.0$, $A_1 = 5.0$, $K_1 = 1.0$, $A_2 = 8.0$, $K_2 = 3.7$, $\kappa = 10.0$, and $M = 1.0$.

#### S3.1. Movie-S1

*Movie-S1*: Time evolution of phase separation in the Tau–tannic acid system at mean protein volume fraction $\bar{\phi} = 0.5$. Starting from a homogeneous state, concentration fluctuations grow into interconnected protein-rich and protein-poor domains that subsequently coarsen and develop sharp interfaces, characteristic of a stable phase-separated state.

##### S3.2. Movie-S2

*Movie-S2*: Time evolution of phase separation in the Tau–tannic acid system at mean protein volume fraction $\bar{\phi} = 0.3$. Starting from a homogeneous state, concentration fluctuations amplify and evolve into protein-rich droplets dispersed within a solvent-rich background. The droplets grow and coarsen over time, illustrating phase separation in the dilute regime governed by the same underlying thermodynamic mechanism.
